# Oxford Nanopore sequencing in a research-based undergraduate course

**DOI:** 10.1101/227439

**Authors:** Yi Zeng, Christopher H. Martin

**Affiliations:** Department of Biology, University of North Carolina at Chapel Hill, NC, USA

**Keywords:** education, third-gen sequencing, long-read sequencing, pupfish, genomics

## Abstract

**Background:** Nanopore sequencing is a third generation genomic sequencing method that offers real time sequencing of DNA samples. Nanopore sequencing is an excellent tool for teaching because it involves cutting-edge sequencing methods and also helps students to develop a research mindset, where students can learn to identify and resolve problems that arise during an experiment.

**Results:** We, as a group of undergraduate biology students, were able to use nanopore sequencing to analyze a sample of pupfish DNA. We were able to accomplish this without computer science backgrounds and only some basic DNA extraction training. Although there were issues, such as inconsistent results across runs, we found it useful as a research learning experience and an application of the skills we learned.

**Conclusions:** As students, it was exciting to be able to experience this technology first hand and apply what we learned in the classroom. Nanopore sequencing holds potential for DNA sequencing of large fragments in real time. It allows students to be acquainted with novel technologies and the theories behind them. However, as with all new techniques, it does not have the same established support, and when students run into difficulties while using nanopore sequencing, it is often difficult to identify what went wrong.

## Introduction

When I first decided to take a seafood mislabeling class I didn’t expect to be able to test developing third-generation DNA sequencing technology or learn about unique evolutionary adaptations from two new species of pupfish from the Bahamas. My course was aimed towards giving undergraduate biology majors a chance to get hands on experience in a research environment. Oxford nanopore sequencing can be applied in a much broader biological context and even dips into computer programming. So, what exactly is nanopore sequencing and why is it important?

DNA sequencing has come a long way from traditional Sanger sequencing. Although Sanger sequencing is still considered the gold standard for accuracy, it requires gel electrophoresis and the use of ddNTPs to identify the sequences of amplified segments of DNA (Obenrader 2003). Sanger sequencing is incredibly accurate but requires lots of processing and thus the cost of sequencing DNA of even relatively short fragments is very high (Goodwin 2016). Next generation sequencing which replaced traditional methods drastically reduced prices of whole genome sequencing (Goodwin 2016). Although these methods made DNA sequencing more accessible, they are generally less accurate (99%) than methods such as Sanger sequencing which can have accuracies as high as 99.999% (Shendure 2008; Morozova 2008). Another major constraint in next generation, or second generation, sequencing is the short read length (50 – 250 bp on Illumina platforms) which makes genome assembly and alignment very difficult (Goodwin 2016).

We are currently approaching the third generation of genomic sequencing which builds further upon the advancements of second generation sequencing. Third generation sequencing addresses the main weakness of second generation sequencing because many platforms can produce reads over 10 kb to even 100 kb or more (Lee 2016). Third generation sequencing does not require PCR amplification as previous generations have. Due to the read lengths and high accuracies, the current third generation sequencing technologies, Pacific Biosciences (PacBio) Single Molecule Real Time (SMRT) sequencing, Illumina Tru-seq Synthetic Long-Read technology, and Oxford Nanopore Technologies sequencing have been able to fill in previous gaps in genomes (Lee 2016). Of the three, Illumina Tru-Seq Synthetic Long Reads is considered to be the most accurate but has much shorter read lengths, requires more DNA, and is more expensive (Lee 2016). PacBio’s SMRT and Oxford Nanopore Technologies sequencing techniques both have high raw error rates, but with certain algorithm techniques both methods can greatly increase their accuracy to almost 99.999% (Lee 2016). The most recent of third generation technologies, Oxford Nanopore Technologies, is greatly limited in the accuracy of their reads in comparison to other third generation techniques but its strength is in its portability. Oxford Nanopore Technologies’ MinION is a device slightly larger than a USB drive that can be plugged into any modern laptop to provide real-time data. Currently in development is an even more portable version that can be run on a smartphone called the SmidgION.

Nanopore sequencing is a third-generation genomic sequencing technique that was only recently commercialized in 2014. The nanopore is a protein with a pore that is only a nanometer wide (Clarke 2009). This tiny pore only allows single molecules such as individual DNA strands to pass through one at a time. These nanopores are embedded into a sheet that has a current applied through it. The current is extremely sensitive to the molecules that pass through it as each nucleotide creates a unique change in the resistance of the nanopore which allows individual nucleotides to be identified individually as they pass through (Deamer 2016). Nanopore sequencing can read DNA sequences more than 100 kb long and the entire run completes within 48 hours. This potentially makes it a low-cost method to sequence genomes, as each kit (including the USB MinION sequencing machine) costs $1,000 for two samples (Meller 2000). Although currently nanopore sequencing is not as accurate in comparison to other third-generation sequencing techniques, what makes this developing technology so exciting is that it offers a possible tool in the future to sequence entire genomes on any lab bench. In fact, it has already proven to be useful in the identification and genome sequencing of diseases in the field, such as Ebola virus (Quick 2016; Schmidt 2017). However, as the newest of the third-generation sequencing technologies, nanopore technologies are continuing to rapidly improve as more accurate and robust nanopores are being developed (Oxford Nanopore Technologies 2017).

Not much has been done in a classroom setting exploring nanopore sequencing or the techniques involved and thus it is a unique opportunity for undergraduates to explore developing technologies. The portability and ease of preparation makes nanopore sequencing ideal in a classroom, because all that students need to analyze data obtained from prepared samples is a working laptop. Previously, only one upper-level class has been well documented which focused on the analysis of DNA sequences produced from nanopore sequencing to identify unknown samples, as well as unknown human DNA (Zaaijer 2016). However, the students did not actually extract the DNA or prepare the sequencing libraries. Analyzing human DNA also presents ethical concerns and falls under institutional review board policies. However, the application of nanopore sequencing can be developed further by teaching many basic laboratory techniques such as proper pipetting and DNA extraction which are necessary to prepare samples. Thus, nanopore sequencing can be used for *de novo* assemblies in the classroom and to introduce students to important laboratory skills and exciting developments in genomic sequencing.

We chose the nanopore sequencing project as our independent research project topic because of its novelty as a portable option for genomic sequencing. We sequenced a sample of muscle tissue from the molluscivore pupfish, *Cyprinodon brontotheroides*, a species of pupfish endemic to San Salvador Island in the Bahamas (Box 1). The genomes of other species of pupfish have already been sequenced so we would also be able to test whether the DNA sequences we got from nanopore sequencing were accurate enough to identify the pupfish sample. Our goal was to use nanopore sequencing to sequence the pupfish genomic DNA and compare the sequence with known data using the Basic Local Alignment Search Tool (BLAST) by the National Center for Biotechnology Information, both as a test to see the accuracy of nanopore sequencing and to see if we, as undergraduates with very little training beforehand, could even successfully use nanopore sequencing.

#### Box 1: Novelty in Caribbean pupfishes

A young, sympatric radiation of pupfishes is endemic to a single Bahamian island despite widespread gene flow and ecological opportunity across the Caribbean (Richards and Martin 2017; Martin 2016a,b; Martin and Wainwright 2011). This 10 kya radiation contains two novel trophic specialists: a scale-eating pupfish and a molluscivore pupfish in additional to a generalist, unique niches among Cyprinodontiform fishes (Martin and Wainwright 2013a,b; McGirr and Martin 2017; Martin et al. 2016). This paradox enables a rare glimpse at the microevolutionary origins of macroevolution-scale novelty: why did novel trophic specialists evolve on only one island across the Caribbean? This is the central question of my research program: predicting the origins of novelty. **Systems like this are rare because the origins of novel adaptations are rare, but enable the chance to study the evolution of novelty in action, rather than millions of years after the fact**.

## Results

We found that it was entirely possible for undergraduate biology students to run nanopore sequencing. From our successful run we generated a massive amount of raw DNA sequences ranging from smaller segments to our longest read of 170 kb (see Table 1).

**Table 1.**
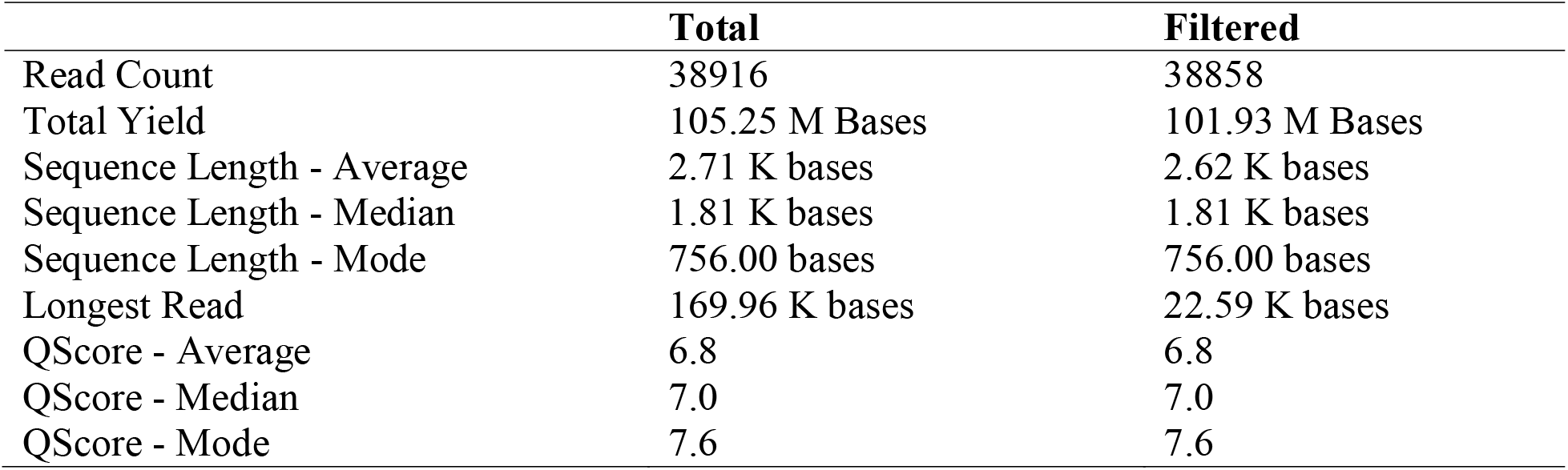
Summary obtained from Metrichor of the run (Oxford Nanopore Technologies 2016). Total results were from all the reads obtained during the run, while filtered results are reads that passed a quality score in MinKNOW. QScore is the quality score assigned to the runs.

We ran a total of three experiments. We recorded the number of active nanopores before conducting each experiment (Table 2). We conducted the quality control experiment using Lambda DNA provided as suggested on the first flow cell before using sample DNA. Our second experimental run using extracted pupfish DNA was successful and the results are reported in Figure 2 and 3.

**Table 2:**
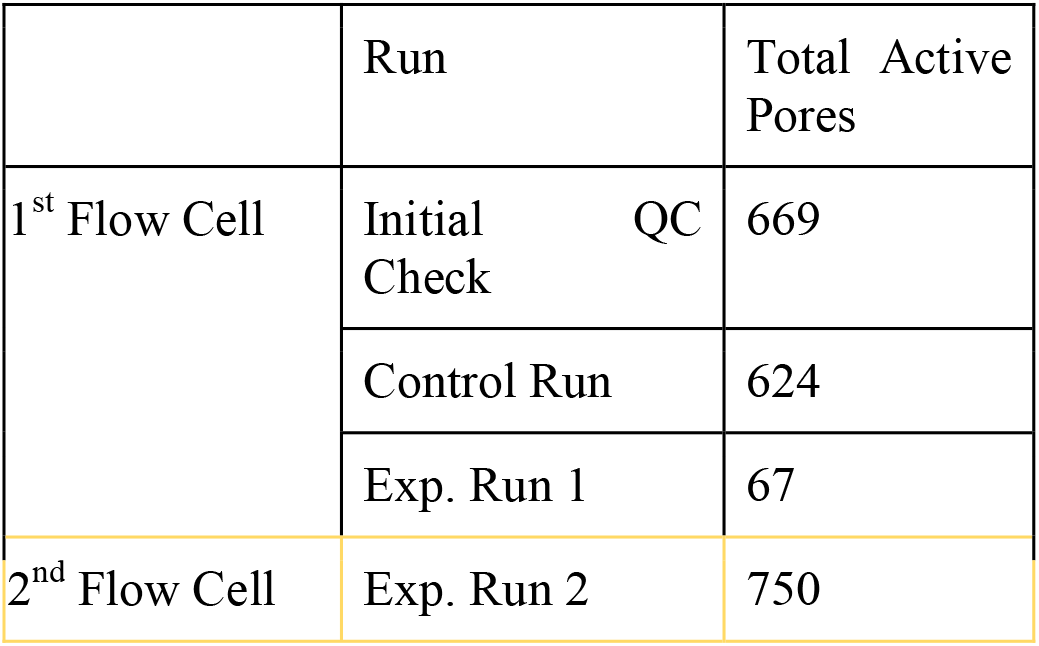
Active Pores Across Runs

**Figure 1.**
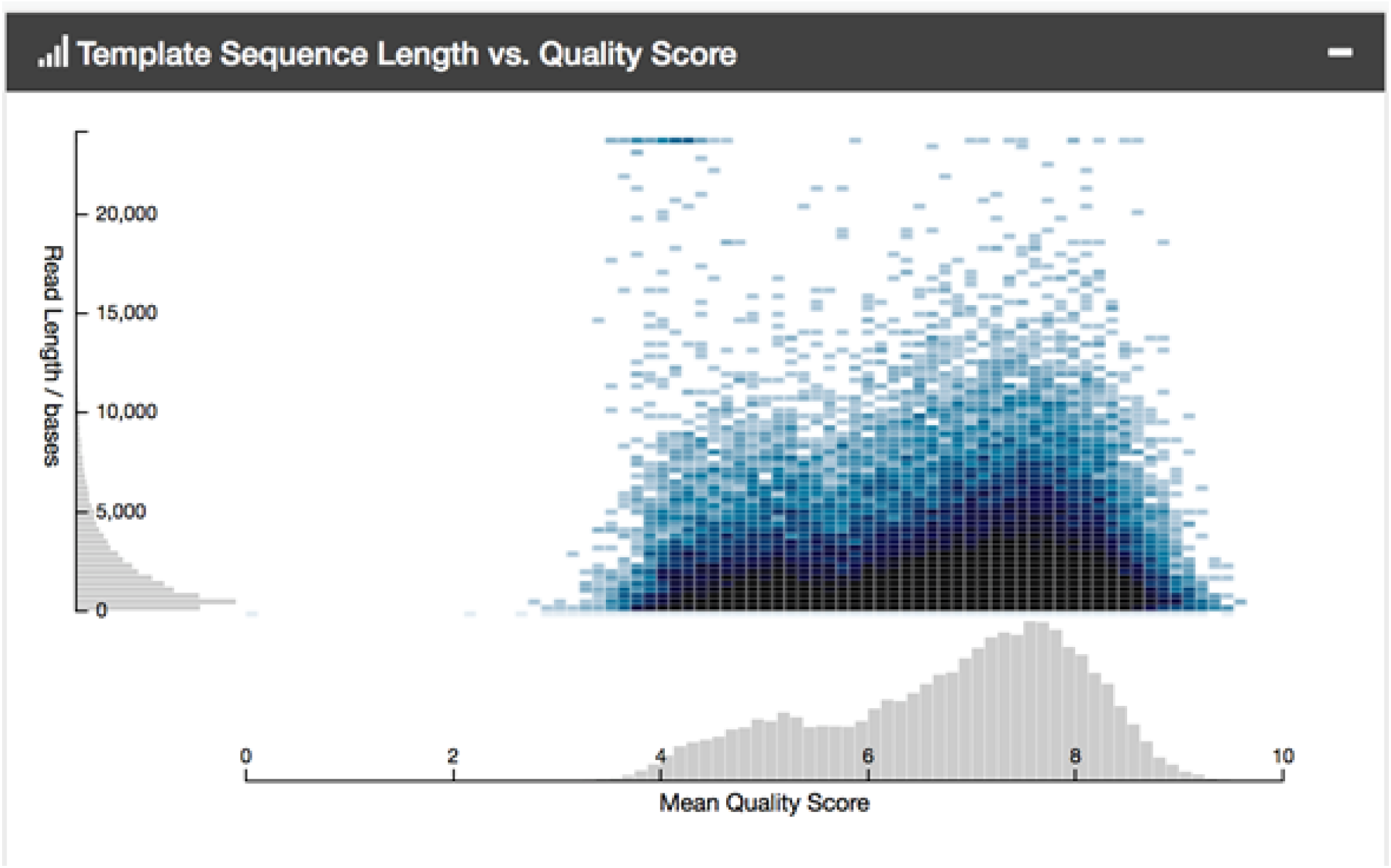
DNA sequence length versus accuracy of the read.

**Figure 2.**
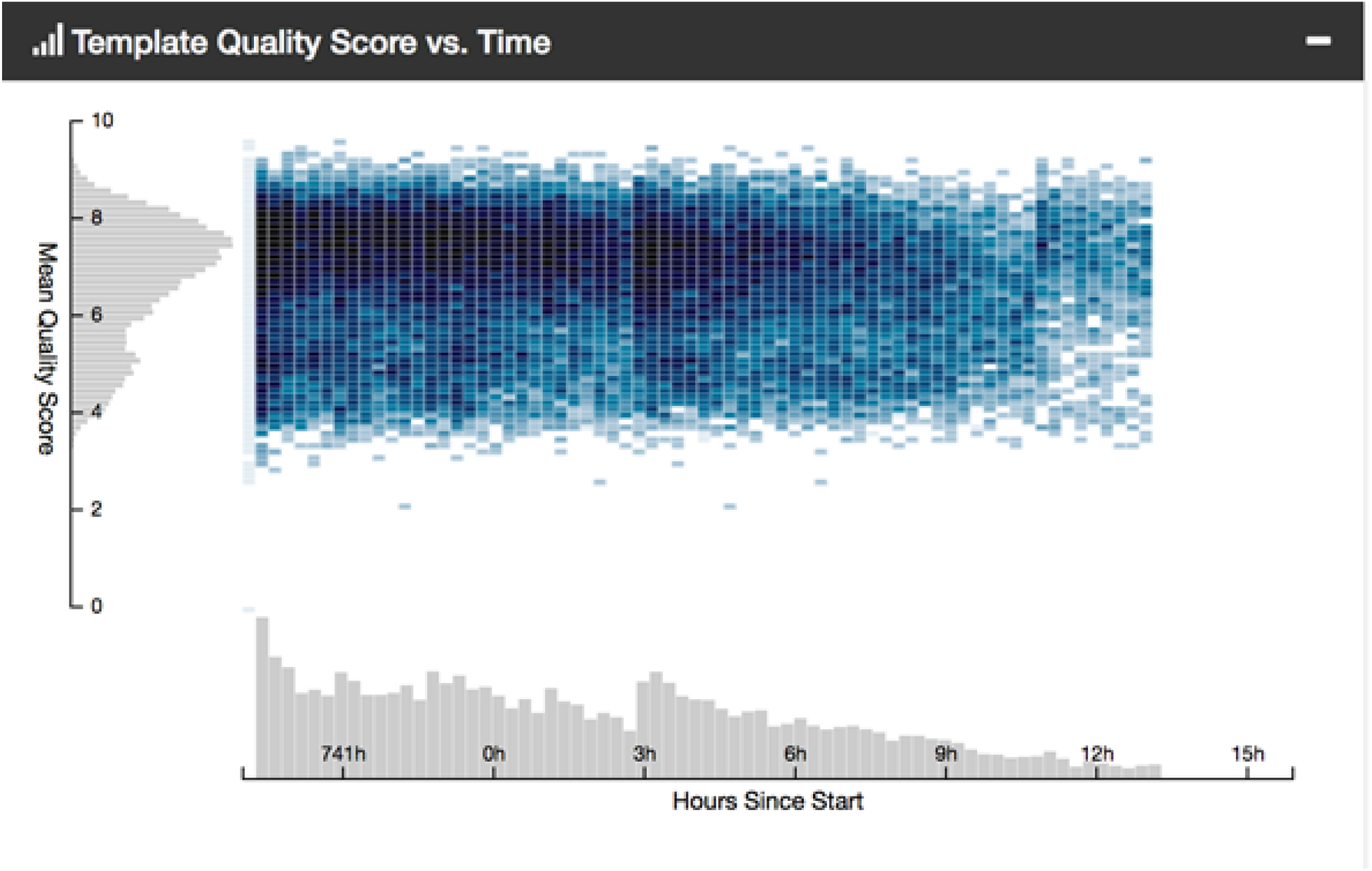
Accuracy of the read versus run time.

**Table 3:** Organisms identified from 10 random sequences. Sample 2 is of a transposon from the green spotted pufferfish. Sample 4 is a guppy, but has a very high e-value.

| Sample | Organisim | E-Value |
| --- | --- | --- |
| 1 | No Match | N/A |
| 2 | Tetraodon nigroviridis transposon Titof2_Tet | 0.0 |
| 3 | No Match | N/A |
| 4 | PREDICTED: Poecilia reticulata spectrin beta chain, non-erythrocytic 4-like (LOC103476183), transcript variant XD, mRNA | 1.7 |
| 5 | N/A | N/A |
| 6 | Populus trichocarpa clone POP002-A07, complete sequence | 0.41 |
| 7 | PREDICTED: Zea mays auxin response factor 17 (LOC103629639), mRNA | 1.3 |
| 8 | Ralstonia insidiosa strain ATCC 49129 chromosome 1, complete sequence | 0.46 |
| 9 | Syphacia muris genome assembly S_muris_Valencia ,scaffold SMUV_scaffold0001048 | 0.029 |
| 10 | Ustilago maydis 521 hypothetical protein partial mRNA | 0.17 |

Metrichor has an application called What’s in my pot (WIMP) that is intended as the analytical component of nanopore sequencing. Currently WIMP can identify bacteria and other unicellular organisms by comparing sequences generated by runs with known genomic sequences and Oxford Nanopore is working to include support for multicellular organisms in the future. WIMP identified *Mycobacterium rhodesiae* and *Delftia* with high classification scores indicating Metrichor’s level of confidence in the identification of the species similar to e-values used by NCBI’s BLAST. Of the two bacteria identified *Mycobacterium rhodesiae*, which had the highest classification score, was very interesting as the pupfish population from which the sample was taken from was contaminated with a mycobacterium infection relatively recently.

Unfortunately, as WIMP does not currently have support for multicellular organisms we had to access the raw data from the run using HDFView-2.13.0 and compare individual sequences through BLAST (National Center for Biotechnology Information). From ten individual sequences we chose at random gathered from BLAST green spotted pufferfish transposons were identified with high accuracy with an e-value of 0.0. Although these are not pupfish, the fish genome is highly conserved between species and the low accuracy of individual reads means that exact matches to the pupfish genome are rare. Furthermore, we analyzed one hundred sequences that passed MinKNOW’s quality filter indicating the program’s confidence in the accuracy of the nucleotides identified (See Appendix 1). Twenty four of the sequences matched a fish species within GenBank, indicating that approximately 24% of reads successfully came from our sample DNA.

## Discussion

My fellow students expressed great interest in nanopore sequencing and took initiative to learn how to use the MinION. It was an opportunity to apply the basic techniques we learned in the class, such as DNA extraction, to nanopore sequencing. Although, as undergraduates, we had no problem prepping and running the library with what was provided with the MinION, we did find multiple problems while running the experiment.

We had difficulties with some of the basics such as opening the files, which are in .fast5 format. HDFView by the HDF Group is a program that can open the files and will present the sequences in FASTA format which can be easily run through BLAST. Our main obstacle, however, was sorting through the raw data and identifying which sequences to BLAST. With computer science backgrounds, it’s possible to tackle this problem, but as none of us had much experience with programming we had to look for alternatives. Beyond Metrichor, the MinION has a lot of user made programs, each with their own pros and cons, most notably Poretools (an extensive list by Next Gen Seek can be found in the references). Poretools is a toolkit that can be used to go through the large amount of raw data and sort the reads for easier analysis (Loman 2014). This allows users who would like to view high quality sequences to access them quickly.

Metrichor, the analytical component of the MinION, can currently only recognize organisms such as bacteria. In identification of pupfish this makes the results from Metrichor less useful as a fish is a multicellular organism. We found it difficult to select high quality reads from the data we generated to BLAST as nanopore sequencing created such large amounts of data and currently has very little user friendly supporting programs for analysis. Luckily, currently Oxford Nanopore is working to continue developing Metrichor so it can search for more organisms from the NCBI database which would resolve the current issue of identifying species from the data obtained. In the end, the easiest method for us was to use HDFView to open the files before using BLAST for individual sequences.

Another problem was that MinION flow cells are advertised to be able to run about three experiments each but active nanopores degrade rapidly, especially after the long runs needed for the experiments. In reality each flow cell can only handle one run, before too many nanopores become degraded. Originally, we had hoped to run more experimental runs, but the experience of the first failed runs taught us how to troubleshoot and improve our methods for the future as we had to go back and identify possible mistakes in our procedure that may have negatively impacted the accuracy of our results. We learned how to identify possible problems that occurred during the experiment and how to avoid those factors in the final experimental run.

Nanopore sequencing can be an incredibly useful educational tool. It brings a portable, user-friendly technology to students and can introduce students of various backgrounds and experience to modern molecular biology. Courses dedicated to exploring nanopore sequencing as a tool for genomic sequencing have many options to choose from. Undergraduate classes can choose to focus on bacterial genomes until WIMP is further updated to include other taxa. Nanopore sequencing can also be used to identify unknown samples and introduce students to how to use tools such as BLAST for identifying organisms from nucleotide sequences. This technology is not necessarily even limited to college classrooms, as there have already been workshops promoting interest in science for young girls that use similar portable DNA identification technologies to great effect (Chacon-Heszele 2016). The ease of access nanopore sequencing offers to the field is also its strength for education as it provides students with an opportunity to witness easier to access and use technologies.

There have been other courses that have also explored aspects of nanopore sequencing as well. Most notably, Columbia University offered a 13-week course that introduced students to the MinION in hackathon sessions which allowed students to generate DNA sequences and use them to identify unknown species or human DNA (Zaaijer 2016). Another example occurred near Acadia National Park in Maine, where researchers trained high school students during the summer to help them sequence samples collected from the park (Krol 2015). There are various ways in which nanopore sequencing can be used in an educational context, and with its lower cost in comparison to other genetic sequencing alternatives and mobile capability it opens the doors for students to experience sequencing on the field.

### Possible lesson plans

#### WIMP

Metrichor’s What’s In My Pot application offers a large variety of options students can explore. One activity students can do is to take environmental swabs of various locations and identify which organisms they find in their samples. Students can choose locations that interest them, such as public transportation, bathrooms, or water from local streams. They can take their samples and purify and extract DNA. Using nanopore sequencing to read their DNA they can then analyze and identify which bacterial species they found in their samples using WIMP. However, until WIMP’s database is updated, the species that can be identified will be limited to bacterial and fungal.

#### Identification of an unknown sample

Students can also use nanopore sequencing as a tool to determine the identity of an unknown sample. Students may be interested in identifying the ingredients of the food they normally consume or teachers could set up a more structured lab for students to identify various samples. For example, if students were interested in figuring out which type of fish were used in their sushi, they could use nanopore sequencing to identify the extracted DNA from a slice of sashimi. Alternatively, teachers could prepare samples of DNA and have students identify which unknown organism they were given.

#### De novo genome assembly

As a third generation sequencing technology, nanopore sequencing is also entirely capable of generating incredibly long reads that are necessary for genome assembly. This approach would require some experience with genome assembly. Students could use results from nanopore sequencing and other sequencing methods to assemble the genome of a sample.

## Conclusion

We found nanopore sequencing to be an incredibly interesting path for our independent project and found that even with very little guidance beyond what is provided online that we could properly run the experiments. Although some basic knowledge of laboratory techniques is necessary, such as pipetting procedures and DNA extraction, the procedure itself for running nanopore sequencing is very straight forward. As we had very little computer science background, our main issue was actually analyzing the data we obtained.

## Methods

### DNA Extraction

The DNA sample was extracted using instructions provided in Qiagen’s DNeasy Blood and Tissue Kit adapted to increase higher yields of long segments of DNA

#### Materials

- Tissue sample
- 1.5 mL microcentrifuge tubes
- Centrifuge
- Buffer ATL
- Proteinase K
- Buffer AL
- 100% ethanol
- DNeasy Mini spin column and 2 mL collection tube
- Buffer AW1
- Buffer AW2
- Nuclease-free PCR water

1. Cut two pupfish filets into smaller pieces and add 180 μL Buffer ATL and 20 μL proteinase K.
2. Mix by inverting instead of vortexing to prevent DNA from fragmenting. Then incubate at 56 □ for 3 hours while periodically mixing the solution about every 15 minutes.
3. At the end of incubation period add 200 μL Buffer AL and incubate for another 10 minutes at 56□
4. Add 200 μL of 100% ethanol to extract the DNA and then pipet the solution into a DNeasy Mini spin column placed in a 2 mL collection tube. Centrifuge at 8000 rpm for one minute and discard the flow through.
5. Place the spin column into a new 2 mL collection tube and add 700 Buffer AW1 for the first wash. Repeat the centrifugation step at 8000 rpm for five minutes.
6. Place the spin column into another new 2 mL collection tube and wash with 700 μL Buffer AW2. Centrifuge at 14,000 rpm for ten minutes. Repeat this step.
7. Elute the DNA by adding 200 μL of Nuclease-free PCR grade water heated up to 55 □ to the center of the spin column membrane. Incubate for 1 minute at room temperature then centrifuge at 8000 rpm for one minute. Repeat this step to increase DNA yield.

### Quality Check of Extracted DNA

Analyze the samples of extracted DNA from pupfish filets to make sure they meet minimum quality requirements for nanopore analysis.

a. Use1 μL of the samples for nanodrop analysis. Check to make sure that the purity is at least 1.8 from Nanodrop OD 260/280 and 2.0-2.2 from Nanodrop OD 260/230.
b. Prepare a 1% agarose gel mixture by adding 1 g of agarose powder for every 100 mL of TAE mixture. Mix and microwave at 1 minute intervals for a total of 3 minutes. Allow the gel mixture to cool and then pour into a gel mold and insert a gel comb. Wait for 30 minutes to let gel solidify. Mix 2 μL of the sample and 2 μL of ethidium bromide dye. Load 2 μL of the sample and a DNA ladder into the gel Check that the average size is greater than 30 kb.
c. Analyze the sample by Qubit by setting up array tubes for two standards and for each of the samples. Prepare the standards by adding 190 μL of Qubit buffer and 10 μL of either standard 1 or 2. Prepare the samples by adding 198 μL of Qubit buffer and 2 μL of sample. vortex the solutions for ten seconds before incubating at room temperature for two minutes. Calibrate the machine with the standards before inserting samples. Dilute the sample to 200 ng per 7.5 μL by using the concentration measured by Qubit.

### MinION Rapid Sequencing Kit

The method was adapted from Oxford Nanopore Technologies protocol for the Rapid Sequencing of genomic DNA for the MinION.

#### Materials

- 1.5 mL microcentrifuge tubes
- 0.2 mL PCR tubes
- Centrifuge
- MinION
- MinION flow cell
- Computer
- 200 ng high molecular weight DNA
- Lambda control DNA
- Nuclease free water
- FRM
- RAD
- NEB Blunt/TA Ligase Master Mix
- RBF-1

1. Prepare the library in the concentrations as shown in Table 1.
2. Incubate at 30°C for one minute then at 75°C for one minute and spin down in a centrifuge.
3. Add 1 μL RAD, the provided adapter and 0.5 μL Blunt/TA Ligase Master Mix and incubate for 5 minutes at room temperature. Place the solution on ice until the prepared library is ready to be loaded into the MinION flow cell. This step fragments the DNA and adds an adapter that can be recognized by the nanopores on the MinION flow cell.
4. Assemble the MinION by inserting the MinION flow cell into the MinION and preform a QC run to check the number of active pores.
5. Prepare the priming buffer by adding 500 μL RBF1 and 500 μL of nuclease free water into a 1.5 mL micro centrifuge tube. Draw back a few μLs of buffer from the sample port to remove air bubbles. Load the priming buffer in ten minute intervals into the sample port.
6. Prepare the library for loading by adding 38 μL of RBF1 and 32 μL of nuclease free water at room temperature into a 1.5 mL micro centrifuge tube. Add 6 μL adapted and tethered library and spin down.
7. Load 75 μL of the sample into the flow cell one drop at a time through the SpotON port under the activator.
8. Load the program through MinKNOW, a software provided for the MinION. MinKNOW should be able to provide data in real time before it can be uploaded and analyzed using Metrichor. Metrichor can check the quality of the DNA sequencing and currently can provide identification of unicellular organisms in the solution.
9. Use the Basic Local Alignment Search Tool (BLAST) by the National Center for Biotechnology Information to compare raw DNA sequences collected by MinION to known genome sequences if necessary. The raw data may be accessed from the run by opening the folder that MinKNOW generates for the run. To manually view the files, which are in .fast5 format, will require a program such as HDFView to open and convert into FASTA format.^20^ The nucleotide sequence in FASTA format can be imported or copied into BLAST and the sequence will be searched against the database. There are alternative methods such as using a user-created program called Poretools which can sort through the raw data and present it in a readable format. However, as it requires some computer science background to be able to use it, our group took 10 random files from the database generated and used HDFView to open the sequences. These 10 random files were then individually entered into BLAST and their results recorded.

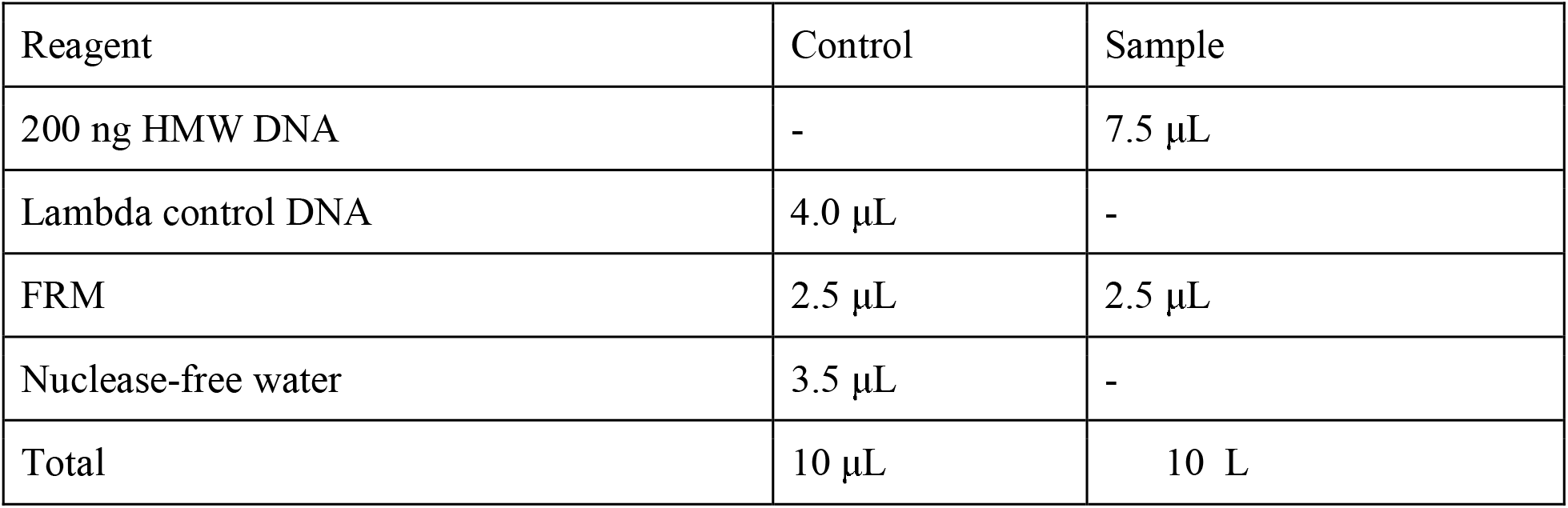

| Reagent | Control | Sample |
| --- | --- | --- |
| 200 ng HMW DNA | - | 7.5 µL |
| Lambda control DNA | 4.0 µL | - |
| FRM | 2.5 µL | 2.5 µL |
| Nuclease-free water | 3.5 µL | - |
| Total | 10 µL | 10 L |

## Ethics approval and consent to participate

Not applicable.

## Consent for publication

All authors have consented to submitting this version of the manuscript for publication.

## Availability of data and material

Raw sequence reads resulting from Oxford Nanopore runs will be uploaded to NCBI’s Short Read Archive. Sequences used for BLAST searches will be provided as supplemental data.

## Competing interests

The authors declare that they have no competing interests.

## Author contributions

YZ prepared genomic libraries, ran the sequencing machine, wrote the manuscript, and analyzed the data. CHM provided funding for the study and contributed to revision of the manuscript.

## Acknowledgements

University of North Carolina at Chapel Hill’s Quality Enhancement Program provided funding for this research to improve student learning outcomes in a research-based undergraduate course (CURE). We thank co-instructor J Bruno for his openness to including nanopore sequencing within the scope of his seafood forensics lab course. We thank YZ’s classmates for their contributions to this project. Permission to collect the pupfish specimen was provided by the Bahamas BEST commission and all animal procedures (breeding and euthanasia) followed approved animal care protocol 15-179.0 from UNC’s Animal Care and Use Committee.

## Appendix 1

During the seafood forensics course students were first taught how to use pipettes to transfer small amounts of liquid and how to extract DNA from raw fish samples using Qiagen DNeasy Blood and Tissue Kit Quick-Start Protocol. Afterwards we checked to make sure the DNA met the quality requirements of Oxford Nanopore Technologies Rapid Sequencing Kit using nanodrop to check the quality and quantity and gel electrophoresis to insure the fragments were mostly over 30 kb.

In total, we had time to run three experiments. The MinION comes with two flow cells, supposedly capable of running three samples each for a total of six experiments. For the first flow cell we completed two runs, one of a control DNA and one of our samples. The first run we did was the suggested control run using lambda DNA provided in the kit for the MinION that acquainted us with how to prepare and tagment DNA and how to prepare that library for loading into the machine. We also tried an experimental run but while running quality control counts of active nanopores on the flow cells we found that the most of the pores had degraded. As expected our first experimental run using pupfish DNA failed, however interestingly enough the control run also failed.

MinKNOW can provide data in real time although the sample runs take about 48 hours to complete. Within the first half hour there was already a lot of data displayed on the automatically updating charts. MinKNOW is a fairly straight forward program where all the user has to do is enter the flow-cell id, an identifier for the experiment run, and to choose whether they are running the control experiment or a sample experiment. The data is also updated to Metrichor where it can be analyzed.

The data generated from Metrichor indicates that all the reads collected from both our control and the first experimental run failed minimum quality filters that indicate how confident the program is in the accuracy of the sequence reads. We attempted to troubleshoot what may have gone wrong. After carefully rereading all the minimum quality requirements for the extracted DNA, we went back and double-checked the quality of the pupfish DNA using Qubit. Here we noticed that we did not dilute the experimental DNA to the proper concentration. After diluting the sample to the proper concentration, we decided to use a fresh flow cell as the drastic decrease in active pores in the previous experimental run indicated less nanopore activity. However, as the control run also failed to meet minimum quality filters, we decided to also increase the attention to detail in preparation of the library for loading into the flow cell. We took extra care with pipetting small amounts of reagents and also used a centrifuge to insure the solution was completely mixed. This run ended up successful and generated massive amounts of data. Metrichor can track the quality of the reads and record the length of each sequence over time

We also needed to use BLAST to identify random sequences we pulled from the folder MinKNOW indicated passed minimum quality filters to see what we could identify as Metrichor could not identify fish yet. HDFView is a free program that can open the fast5 format MinKNOW saves the sequencing reads in so we used it to analyze a hundred sequences. Then we took the nucleotide sequences and used NCBI’s BLAST tool to see what matched up and the quality of the matches (See Appendix 1).

**Table 3:**
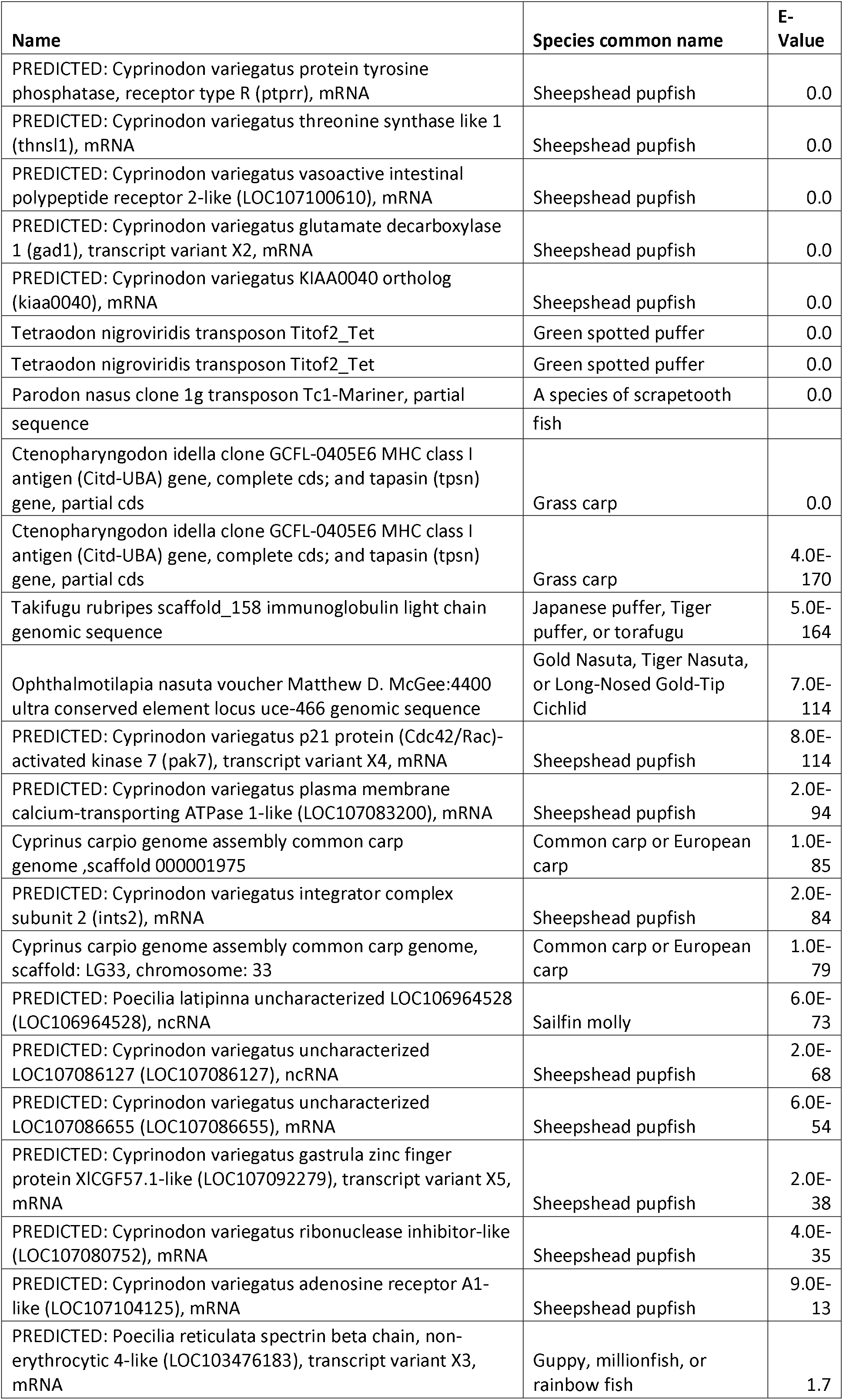
Identified fish sequences from one hundred random successful reads generated using nanopore sequencing.

Out of one hundred samples searched, we found 24 reads matched any fish species. The most commonly identified species was that of the sheepshead pupfish, *Cyprinodon variegatus*, closely related to our sample tissue, *Cyprinodon brontotheroides* (which does not have any GenBank entries). Various bacteria were identified but with very high e-values. There was a single outlier of a sequence that identified as a wild boar that had an e-value of 4.0E-37.

